# Isolated *BAP1* loss in malignant pleural mesothelioma predicts immunogenicity with implications for immunotherapeutic response

**DOI:** 10.1101/2022.05.06.490947

**Authors:** Hatice Ulku Osmanbeyoglu, Drake Palmer, April Sagan, Eleonora Sementino, Joseph R Testa

## Abstract

Malignant pleural mesothelioma (MPM), an aggressive cancer of the mesothelial cells lining the pleural cavity, lacks effective treatments. Multiple somatic mutations and copy number losses in tumor suppressor genes (TSGs) *BAP1, CDKN2A/B*, and *NF2* are associated with MPM. The impact of single versus multiple losses of TSG on MPM biology, the immune tumor microenvironment, clinical outcomes, and treatment responses are unknown. Tumors with alterations in *BAP1* alone were associated with a longer overall patient survival rate compared to tumors with *CDKN2A/B* and/or *NF2* alterations with or without *BAP1* and formed a distinct immunogenic subtype with altered transcription factor and pathway activity patterns. *CDKN2A/B* loss consistently contributed to an adverse clinical outcome. Since the loss of only *BAP1* was associated with the PD-1 therapy response signature and higher *LAG3* and *VISTA* gene expression, it is a candidate for immune checkpoint blockade therapy. Our results on the impact of TSG genotypes on MPM and the correlations between TSG alterations and molecular pathways provide a foundation for developing MPM therapies.

## Introduction

Malignant pleural mesothelioma (MPM) is a rare, devastating cancer of the lining of the lung and thoracic cavity with a 5-year survival rate of < 10%. About 70% of MPM cases are associated with occupational or environmental exposure to asbestos fibers^1,2^. MPM is broadly divided into three histological subtypes with varying biological and clinical behaviors: epithelioid (median survival of ~14.5 months), sarcomatoid (5.5 months) and biphasic (9.5 months), the latter being a combination of the former types^3^. The incidence of MPM has continued to increase in many parts of the world,^4^ and it is predicted to increase dramatically in certain developing countries such as India^5^, where the use of asbestos has increased exponentially with few precautions taken. Moreover, carbon nanotubes, often used in manufacturing and for sportswear, induce inflammation as well as a cellular injury response similar to the *in vitro* response to asbestos fibers^6^, which might lead to a future increase in MPM incidence^7^. The standard of care for MPM with pemetrexed platinum-based chemotherapy extends survival by only 2–3 months^8^. A combination of nivolumab and ipilimumab, the first drug regimen approved by the FDA for MPM since 2004, produced a four-month improvement in overall survival of MPM patients compared with those receiving cisplatin or carboplatin plus pemetrexed^9^. However, only a minority of MPM patients responded to immune checkpoint therapy, and the reasons why most patients did not respond are poorly understood. The failure of clinical trials and a dearth of effective treatments may be due to our poor understanding of this disease and the lack of molecular biomarkers for patient selection.

Loss-of-function mutations and copy number losses in tumor suppressor genes (TSGs) are common in MPM patients, whereas activating mutations of proto-oncogenes are rare^10–13^. The most common losses of TSGs in the genomes of human MPM tumors are in *BAP1* (25%-60% of cases), which encodes a deubiquitinating enzyme originally identified as a BRCA1 interacting protein; *CDKN2A/B* (40%-45% of cases), which encodes the cell cycle inhibitors p16INK4A/p14ARF and p15INK4B, respectively; and *NF2* (20%-50% of cases), which encodes the cytoskeletal scaffolding protein Merlin^10–12^. Although loss or inactivation of these TSGs occur in combination in ~35% of MPM cases, a recent evolutionary analysis by Zhang et al.^13^ suggests that *BAP1* loss occurs early in the evolution of MPM, whereas *NF2* loss occurs later in disease progression. Among the TSGs commonly implicated in MPM, only *CDKN2A/B* has been associated with poor survival^10^. Since it is difficult to develop therapies that target TSG alterations, little progress has been made in a gene-targeted approach to MPM treatment.

The impact of single gene versus multiple TSG losses on MPM biology, patient outcome, or treatment response is largely unknown. To address these gaps in our knowledge, we examined how single gene loss of *BAP1, NF2*, or *CDKN2A* versus multiple TSG gene losses affected clinical outcomes and response to therapy in MPM. We show herein that loss of *BAP1* only is associated with a better outcome and a PD-1 therapy response signature and, thus, is a candidate for immune checkpoint blockade therapies. We also provide evidence for altered transcription factors and pathways associated with each TSG/TSG combination genotype.

## Results

### Association of patient survival with TSG genotypic groups in MPM

Point mutations in *BAP1, NF2*, and *CDKN2A* comprised 28%, 28% and 26%, respectively, of all mutated genes in a dataset of ten large MPM whole-genome or targeted screens^10,12,14–21^ from the Catalog of Somatic Mutations in Cancer database^22^. To determine whether losses in each of these three genes or gene combinations could better stratify clinical outcomes, we analyzed MPM datasets from The Cancer Genome Atlas (TCGA)^10^ (n = 86) and Memorial Sloan Kettering-Integrated Mutation Profiling of Actionable Cancer Targets (MSK-IMPACT, targeted screen)^23^ (n = 61) and divided patients into eight groups: *BAP1* loss alone (B-; TCGA: n = 11, ~13%; MSK-IMPACT: n = 20, 33%), *NF2* loss alone (N-; TCGA: n = 7, ~8%; MSK-IMPACT: n = 2, ~3%), *CDKN2A/B* loss alone (C-; TCGA: n = 16, ~18%; MSK-IMPACT: n = 5, 8%), combined losses in *CDKN2A/B* and *NF2* (N-C-; TCGA: n = 7, 8%; MSK-IMPACT: n = 3, 5%), losses in *BAP1* and *CDKN2A/B* (B-N-; TCGA: n = 3, 3%; MSK-IMPACT: n = 6, ~10%), losses in *BAP1* and *NF2* (B-C-; TCGA: n = 6, 7%; MSK-IMPACT: n = 3, ~5%), losses in *BAP1, NF2*, and *CDKN2A/B* (B-N-C; TCGA: n = 11, ~13%; MSK-IMPACT: n = 2, ~3%), and losses in genes other than these three driver TSGs (B+N+C+; TCGA: n = 26, 30%; MSK-IMPACT: n = 20, 33%) (**Fig. 1A–B**). There are several differences in percentages for the different TSG status between the TCGA and the MSK-IMPACT datasets. The main difference appears to be the considerably higher incidence of alterations in *BAP1* alone seen in the MSK-IMPACT dataset (33%) versus the TCGA dataset (~13%). One reason might be due to the fact that the TCGA data was based on whole exon sequencing whereas MSK-IMPACT was based on a targeted screen, which may have identified a higher percentage of whole exon deletions of *BAP1*. Notably, in early reports, performed with Sanger sequencing, revealed point mutations in 20-25% of sporadic MPMs^14,24^. Subsequent deletion mapping and multiplex ligation-dependent probe amplification studies have identified alterations of BAP1 in 50-60% of MPMs, with the increase due to inactivating deletions of entire *BAP1* exons^25,26^.

**Fig. 1.**
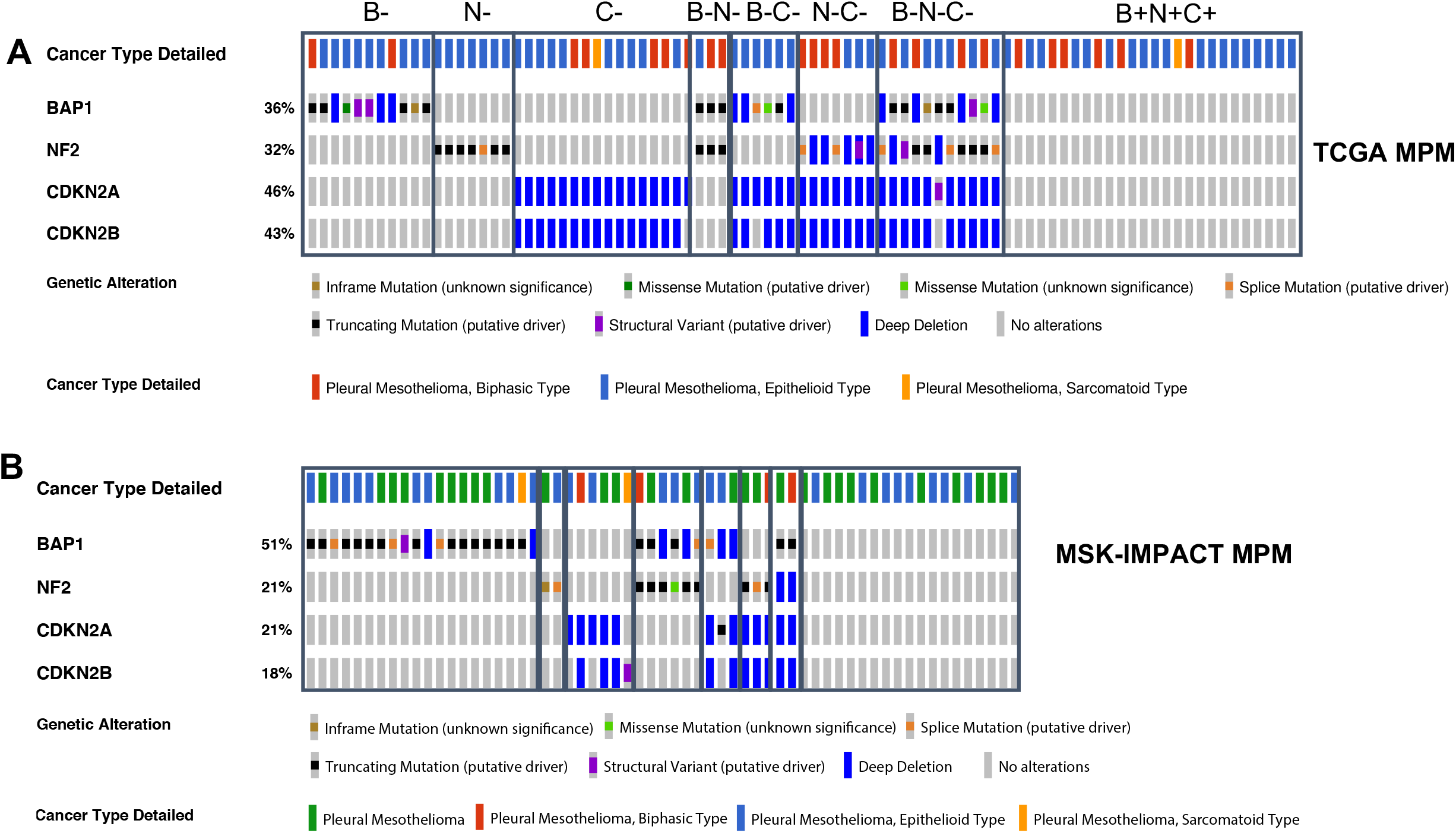
*BAP1, NF2*, and *CDKN2A/B* tumor suppressor gene losses in human malignant pleural mesotheliomas (MPM) **(A)** TCGA and **(B)** MSK-IMPACT cohort. *BAP1* (B), *NF2* (N), and *CDKN2A* (C).

We evaluated the association of overall survival (OS) with tumors with mutations in *BAP1, NF2*, or *CDKN2A* by Kaplan-Meier analysis in the TCGA and MSK-IMPACT MPM datasets. Notably, tumors with a loss of only *BAP1* were significantly associated with longer survival compared with tumors with alterations in *CDKN2A* or/and *NF2* (**Fig. 2A–B**). In contrast, loss of *CDKN2A/B*, with or without loss in *BAP1* or *NF2*, was associated with poor survival, as previously reported^10^.

**Fig. 2.**
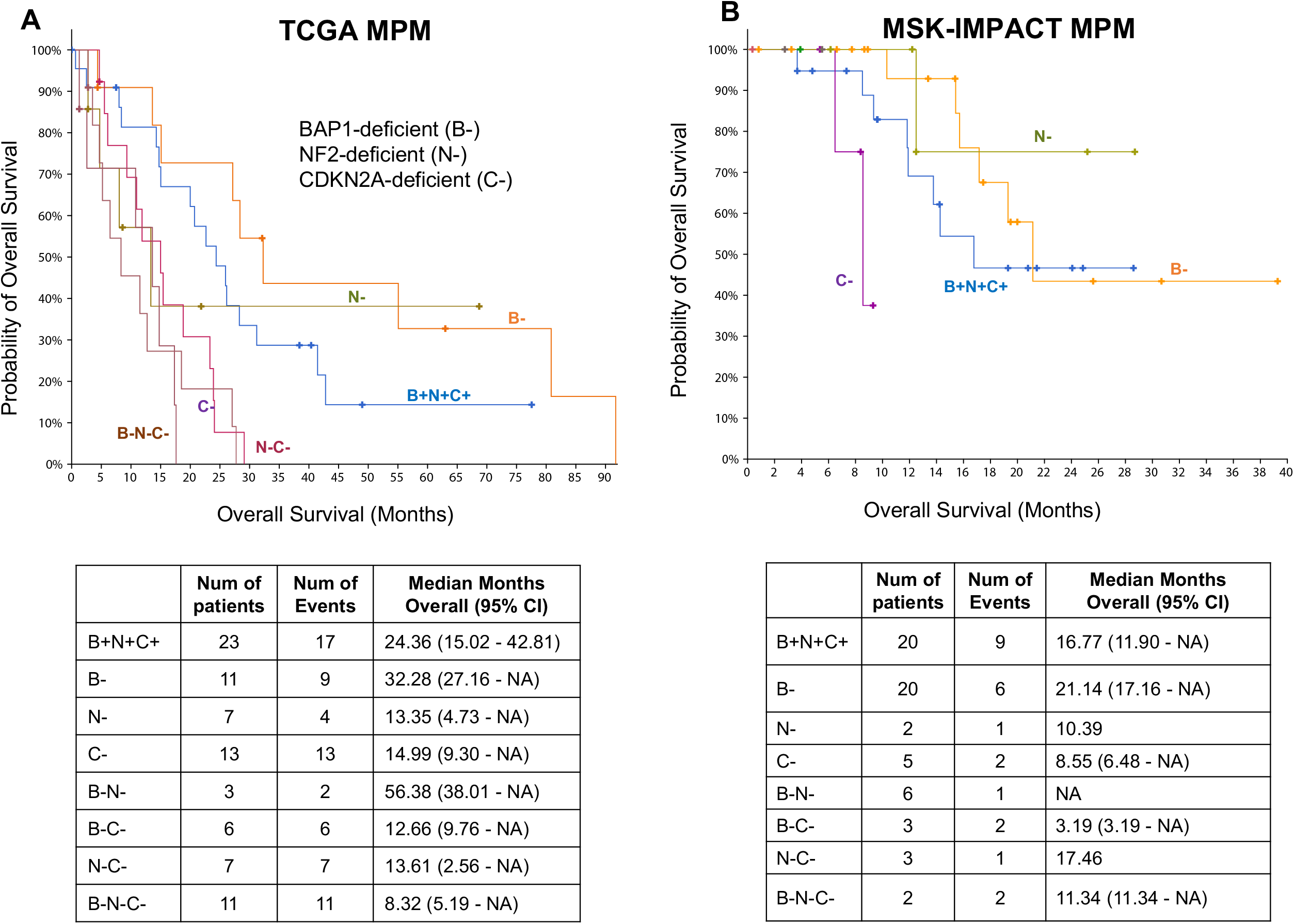
The association of TSG genotypes with overall survival in MPM. Kaplan-Meier plots of clinical outcomes based on *BAP1* (B-), *NF2* (N-), and *CDKN2A* (C-) TSG genotype combinations. B+N+C+ refers to tumors that have alterations of genes other than *BAP1, NF2*, or *CDKN2A/B*. TCGA **(A)** and MCK-IMPACT **(B)** MPM datasets. We filtered groups with less than five samples for survival analysis.

### Association of PD-1 therapy response with TSG genotypic groups in MPM

We performed gene set variation analysis (GSVA)^27^ using published gene expression-based treatment-response signatures to determine differences in patient responses to standard pemetrexed chemotherapy based on TSG genotypes. We found an expression-based signature derived from non-small cell lung cancer that predicted resistance to pemetrexed^28^. To determine differences in sensitivity to cyclin-dependent kinase inhibitors and to drugs that inhibit DNA repair among the eight TSG genotypic groups, we collected a signature of resistance to palbociclib derived from breast cancer^29^, which is shown as GSVA scores for the two signatures across the MPM TSG genotypic groups **(Fig. 3A)**. On average, MPM tumors with *CDKN2A/B* loss, with or without *BAP1* or *NF2* losses, were more resistant to pemetrexed and palbociclib, whereas tumors with only *BAP1* loss or with *BAP1* and *NF2* losses were more sensitive (a lower GSVA score).

**Fig. 3.**
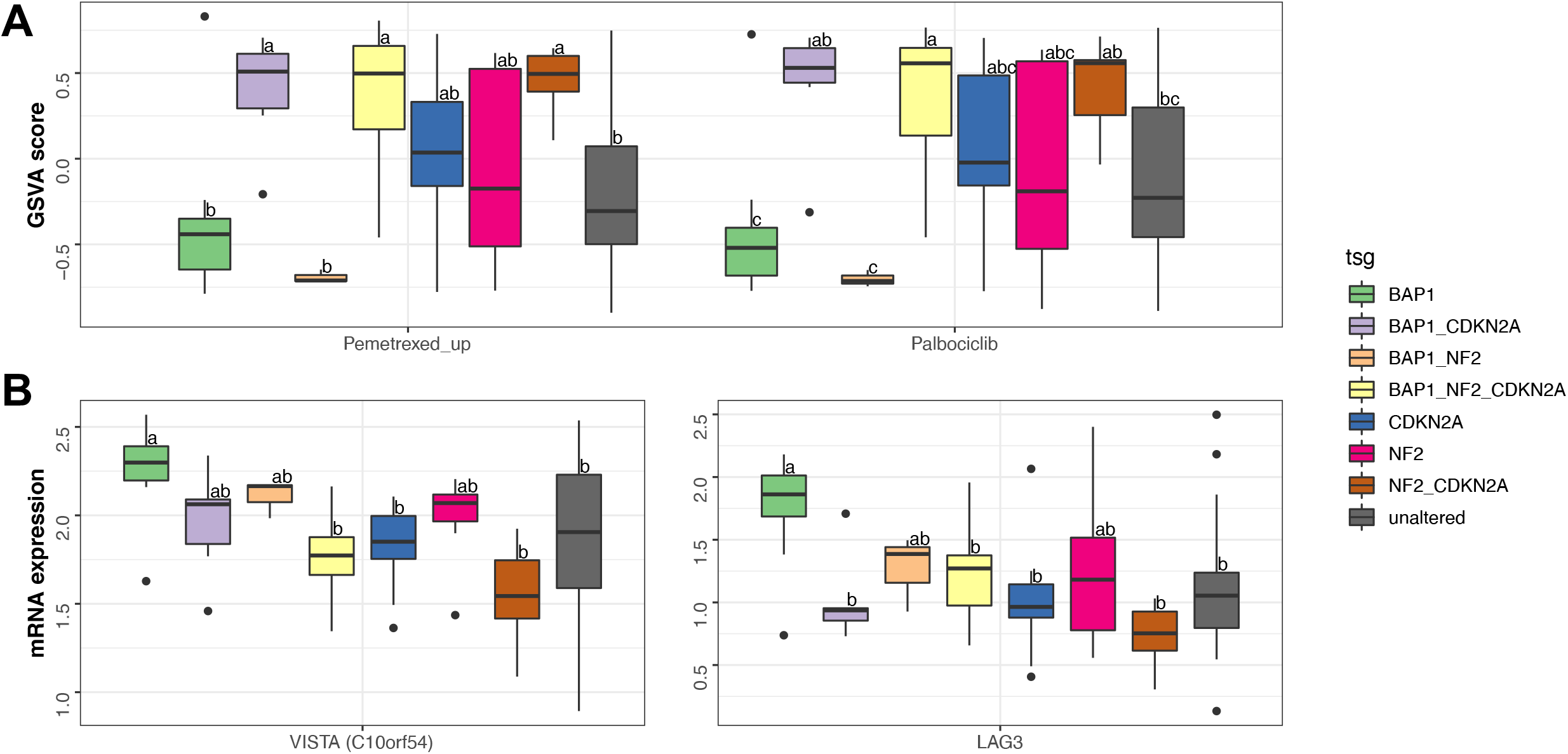
The association of TSG genotypes with drug responses in MPM. **(A)** Enrichment for signatures of resistance to chemotherapy and targeted therapy in patients with MPM (pemetrexed, *left panel*; palbociclib, *right panel*). Positive versus negative GSVA scores (y-axis) indicate upregulation or downregulation of the signature in each patient, respectively. **(B)** Comparison of immune checkpoint mRNA expression levels as a function of TSG genotypes in 86 MPM samples from the TCGA cohort. Samples with different letters exhibited statistically significant mRNA expression or GSVA score differences (ANOVA, Tukey’s HSD, adjusted P-value < 0.1)

To evaluate the connection between TSG genotypes and resistance to PD-1 therapy, we tested a gene expression signature predictive of benefit from immune checkpoint inhibitor (ICI) treatment^30^. Strikingly, we found that tumors with loss of only *BAP1* (B-) were represented only in the anti-PD-1-responsive subgroup (P = 0.0032), whereas N-C-tumors were observed only in the anti-PD-1 resistant subgroup (P = 0.0243) (**Supplementary Table 1**). By comparing immune checkpoint mRNA expression levels as a function of TSG genotypes, we found that expression of *LAG3* and *C10orf54* (*VISTA*) was higher in B-tumors (**Fig. 3B)**, but mRNA levels for other immune checkpoint genes, including *PD-1, PD-L1, TIGIT*, and *CTLA4*, were not associated with TSG genotypes (**Supplementary Figure 1**). In summary, tumors with loss of only *BAP1* formed a distinct MPM subtype that was associated with significantly longer overall patient survival, a better therapeutic response, and higher expression of *LAG3* and *VISTA*.

### Transcription factors and pathways associated with TSG genotypic groups in MPM

To determine whether the eight MPM TSG genotypes impacted similar or distinct molecular pathways, we analyzed transcription factor activity using gene expression data via the Integrated System for Motif Activity Response Analysis (ISMARA)^31^ and sample-specific pathway enrichment analysis based on GSVA^27^. **Fig. 4A** summarizes a set of transcription factors (TFs) (false discovery rate [FDR] < 0.1) and pathways (FDR < 0.05) associated with each TSG group. In particular, C-tumors were associated with increased activity of *MYBL1* (also known as *AMYB*), which is involved in the control of cell survival, proliferation, and differentiation^32^, and *HMGA2*, which encodes a putative transcription factor that belongs to the nonhistone chromosomal high-mobility group protein family. B-N-tumors were associated with increased activity of *TRAP2E*. Consistent with TF activity patterns, tumors with loss of *CDKN2A* were associated with increased activity of pathways involved in the cell cycle and epithelial-mesenchymal transition (EMT) (**Supplementary Table 2**).

**Fig. 4.**
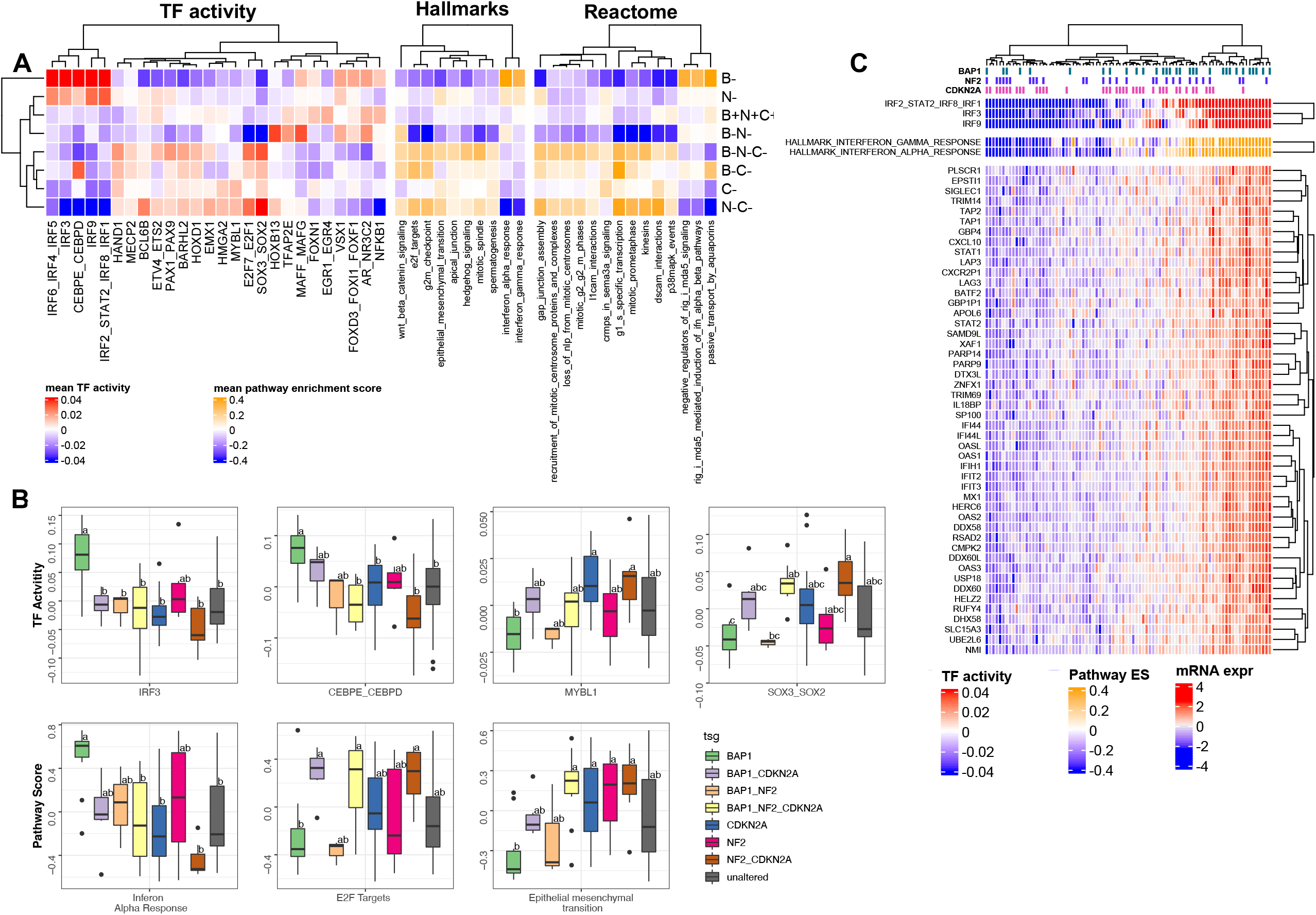
The association of TSG genotypes with TF activity and pathway patterns. (**A)** Heatmap showing the mean inferred TF activity and pathway enrichment scores for each TSG genotype. **(B)** Boxplots indicate the distribution of inferred IRF3, CEBPE_CEBPD, and MYBL1 TF activities and interferon-alpha response E2F targets and epithelial to mesenchymal transition enrichment score (ES) across tumor TSG genotypes. Samples with different letters exhibit statistically significant TF activity or GSVA score (ANOVA, Tukey’s HSD, P < 0.1). **(C)** The top heat map shows tumors clustered by the inferred IRF family TF activities. The middle panel shows the pathway ES for interferon pathways for each tumor based on clustering by TF activities. The bottom panel shows the mRNA expression profile for each tumor for genes highly correlated with IRF TF activity (Pearson correlation > 0.75).

Whereas B-tumors were associated with increased activity of interferon regulatory factors (IRFs) that have a role in immunity, the C/EBP family of TFs has many cellular functions, including proliferation, differentiation, apoptosis, and ER stress. Consistent with the TF activity patterns, B-tumors were associated with increased interferon-alpha and interferon-beta signaling activity. Genes that showed increased expression that correlated with IRF TF activities and pathway score included interferon response genes, such as *IFIT3, OAS2, OAS3, IFIH1, STAT1, DDX60*, and *DDX58*; genes regulating the CGAS-induced type I interferon signaling pathway, such as TRIM14; and transporters associated with antigen processing such as *TAP1* and *TAP2* (**Fig. 4C**). In summary, we identified a group of MPM patients with loss-of-function mutations and/or copy number loss only in *BAP1* that define a distinct molecular subtype associated with a high interferon response.

## Discussion

Precision oncology, which has been successful in treating a variety of cancers, is still in its infancy in MPM. While there are ongoing clinical trials for MPM, there are no defined molecular markers that can be used to predict the efficacy of treatments. Losses of *BAP1, NF2*, and *CDKN2A* TSGs are thought to play a critical role in MPM pathogenesis. In this study, we provide a comprehensive analysis of data on MPM tumors with loss in one or more of these specific TSGs, which are thought to be critical drivers of MPM pathogenesis.

Tumors with loss in only *BAP1* showed a distinct pattern of expression of inflammatory tumor microenvironment genes, including activation of interferon signaling and IRF TFs and high *LAG3* and *VISTA* expression. Interferon production is a defense response that recruits and activates immune cells and has been studied extensively in cancer. Activation of the interferon response in cancer cells^33^, possibly by genomic instability through the cGAS– STING pathway^34^, may affect immune cells in the tumor microenvironment and the tumor response to immunotherapies. In tumors, type I interferons are secreted by cancer cells and dendritic cells (DCs) in response to DNA fragments that activate the cGAS/STING pathway and result in T cell priming and antitumor activity. Thus, loss of *BAP1* may serve as a predictive and prognostic biomarker for MPM to improve disease stratification and therapy. Corroborating our findings, *BAP1* loss has been shown recently to be correlated with perturbed immune signaling in malignant peritoneal mesothelioma^35^. Overall, our results indicate that patients with tumors showing loss in only *BAP1* could be prioritized for immune checkpoint blockade therapies.

## Materials and Methods

### Data and preprocessing

TCGA data were downloaded through the Broad Institute TCGA GDAC firehose tool. RNA-seq data were available for 86 samples. Genetic alteration data (copy number alteration and mutation) for *CDKN2A/B, NF2*, and *BAP1* were retrieved from cBioPortal and clinical data from an online portal for data from the TCGA project and MSK-IMPACT^23^ (http://www.cbioportal.org).

### Survival analyses

OS was defined as the time from diagnosis to death resulting from any cause. Survival curves were estimated using the Kaplan-Meier method and compared with the log-rank test.

### Motif activity analysis

To analyze activities of transcription factor binding motifs (TFBM) from RNA-seq data, we used ISMARA^31^.

### Gene set enrichment analysis

GSVA and single-sample gene set enrichment analysis (ssGSEA) were performed using the GSVA R package (version 1.40.1)^27^ on the pemetrexed and palbociclib response signature and pathway enrichment analysis. For pathway enrichment analysis, we obtained pathway annotations from the Molecular Signatures Database (MsigDB)^36^, a collection of hallmarks of cancer and REACTOME pathways (c2.all.v7.1.symbols.gmt). The log-transformed TMM (trimmed mean of M values) normalized TPM (transcripts per million) counts were used as input to the GSVA package.

### Statistical analysis and visualization

All statistical tests in the exploratory analysis were performed using R version 4.1.1 and associated packages. The statistical analyses for differences in mRNA expression, GSVA score, and TF activity were performed using one-way ANOVA and post hoc Tukey’s HSD (honestly significant difference), with an adjusted p-value cut off of 0.1. One or more letters were assigned to each sample using the multcompView package in R (version: 0.1-8). Assignments were made such that any two samples that had a statistically significant difference did not share any letters.

Graphs were generated using RColor-Brewer (version: 1.1 2), ggplot2 (version: 3.3.3) ComplexHeatmap (version: 2.4.3), ggrepel (version: 0.9.1), and circlize (version: 0.4.13) packages. For general data analysis and manipulation, dplyr (version: 1.0.7), matrixStats (version: 0.59.0) and data.table (version: 1.14.0) were used.

## Acknowledgments

The results published here are, in whole or in part, based on data generated by The Cancer Genome Atlas project established by the NCI and NHGRI (accession number: phs000178.v7p6). Information about TCGA and the investigators and institutions that constitute the TCGA research network can be found at http://cancergenome.nih.gov/. This work was supported by NIH award R00CA207871. This research was supported in part by the University of Pittsburgh Center for Research Computing through the resources provided.

## Author contributions

HUO conceived the study, performed data analysis, and wrote the manuscript. DP performed the ISMARA analysis. AS assisted in statistical analysis. HUO, ES, and JRT analyzed data. JRT helped to write the manuscript.

## The paper explained

### Problem

Malignant Pleural Mesothelioma (MPM) is a highly aggressive, therapy-resistant cancer with a well-established inflammatory etiology and has no cure. Immune checkpoint inhibition (ICI) therapy has shifted treatment paradigms in many types of cancers, and recent clinical data have shown promise for improving MPM treatment. However, response to ICI therapy has been neither uniform nor predictable. The genomic landscape of MPM is primarily characterized by alterations in tumor suppressor genes (TSGs) (~70%), particularly *BAP1, CDKN2A/B* and *NF2*. The impact of an isolated TSG loss versus multiple concurrent TSG losses on clinical outcome, treatment response and MPM biology and the immune tumor microenvironment are unclear.

### Results

Here, we showed the effect of TSG loss combinations on clinical outcome, therapeutic response, and molecular pathways in MPM. Our manuscript presents 3 key findings:

1. Tumors harboring isolated *BAP1* loss have a significantly longer overall survival compared to tumors harboring co-occurrent *CDKN2A* and/or *NF2* loss.
2. Tumors harboring isolated *BAP1* loss form a distinct immunogenic subtype characterized by distinct transcription factor and pathway activity patterns involved in the interferon pathway.
3. Isolated *BAP1* loss is associated with a predicted PD-1 therapy response signature, higher LAG3 and VISTA gene expression and, thus, is a candidate for immune checkpoint blockade therapies.

### Impact

Our study helps fill the knowledge gap between isolated and concurrent TSG loss and their impact in MPM, reveals connections between mutations/loss and molecular pathways, and lays a foundation for developing new therapies for MPM.

## Data availability

RNA-seq gene expression data from TCGA’s Firehose data run (https://confluence.broadinstitute.org/display/GDAC/Dashboard-Stddata). Genetic alteration data (copy number alteration and mutation) were retrieved from cBioPortal and clinical data from an online portal for data from the TCGA project and MSK-IMPACT^23^ (http://www.cbioportal.org).

## Expanded View Figure and Table legends

**Supplementary Figure 1:** Comparison of immune checkpoint gene mRNA expression levels as a function of TSG genotypes in 86 MPM samples from the TCGA cohort.

**Supplementary Table 1:** The anti-PD-1-resistant mRNA signature was used to predict the subgroups. TCGA MPM tumors predicted to be anti-PD-1-sensitive tumors were enriched in samples with *BAP1* loss only. A two-sided proportion test was used to compare each genotype to the B+N+C+ group.

**Supplementary Table 2:** Candidate TF regulators (10% FDR) based on **Fig 4A**. Functional annotations were determined from terms overrepresented from the canonical pathway and from the Gene Ontology ‘Biological Process’ gene sets associated with the candidate regulator based on ISMARA analysis.

