## Supplementary Information for "Isolated *BAP1* loss in malignant pleural mesothelioma predicts immunogenicity with implications for immunotherapeutic response"

### Expanded View

**Supplementary Figure 1:** Comparison of immune checkpoint gene mRNA expression levels as a function of TSG genotypes in 86 MPM samples from the TCGA cohort.

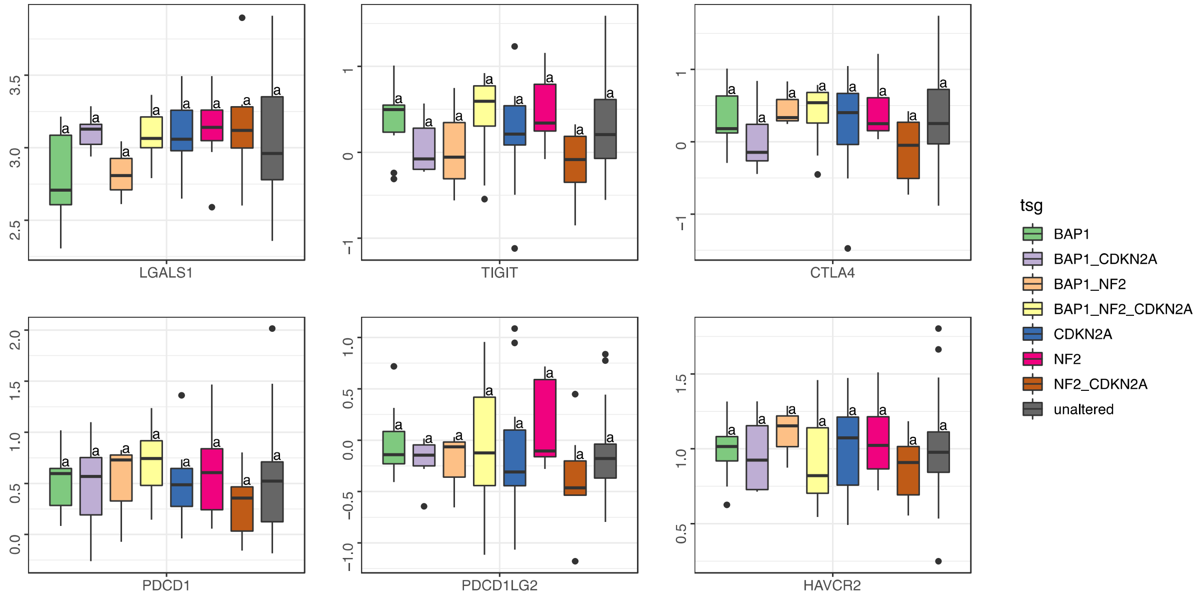

**Supplementary Table 1:** The anti-PD-1-resistant mRNA signature was used to predict the subgroups. TCGA MPM tumors predicted to be anti-PD-1-sensitive tumors were enriched in samples with *BAP1* loss only. A two-sided proportion test was used to compare each genotype to the B+N+C+ group.

| Genotype | Responsive | Resistant | P-value |
|---|---|---|---|
| B+N+C+ | 9 | 10 | 1 |
| B- | 11 | 0 | 0.0032 |
| N- | 4 | 3 | 0.6584 |
| C- | 3 | 9 | 0.2130 |
| B-N- | 0 | 2 | 0.1978 |
| B-C- | 3 | 3 | 0.9104 |
| N-C- | 0 | 7 | 0.0243 |
| B-N-C- | 3 | 6 | - |

**Supplementary Table 2:** Candidate TF regulators (10% FDR) based on **Fig 4A**. Functional annotations were determined from terms overrepresented from the canonical pathway and from the Gene Ontology 'Biological Process' gene sets associated with the candidate regulator based on ISMARA analysis.

| Candidate regulator | Ontologies associated with geneset |
|---|---|
| **IRF2_STAT2_IRF8_IRF1** | Type I interferon (alpha/beta IFN) Pathway |
| **IRF6_IRF4_IRF5** | Type I interferon (alpha/beta IFN) Pathway |
| **IRF9** | CXCR3-mediated signaling events, type I interferon (alpha/beta IFN) pathway |
| **IRF3** | Type I interferon (alpha/beta IFN) Pathway |
| **CEBPE_CEBPD** | Endogenous TLR signaling, validated targets of C-MYC transcriptional repression |
| **VSX1** | AP-1 transcription factor network |
| **HOXD1** | Genes involved in interleukin-7 signaling |
| **BARHL2** | Genes involved in NCAM1 interactions |
| **EMX1** | Genes involved in collagen formation |
| **BCL6B** | Syndecan-4-mediated signaling events |
| **ETV4_ETS2** | Genes involved in kinesins |

| | |
|---|---|
| **EGR1_EGR4** | Wnt signaling network |
| **MYBL1** | chondroitin sulfate binding |
| **HMGA2** | Genes involved in regulation of complement cascade |
| **HAND1** | regulation of branching involved in salivary gland morphogenesis by mesenchymal-epithelial signaling |
| **MECP2** | Genes encoding enzymes and their regulators involved in the remodeling of the extracellular matrix |
| **PAX1_PAX9** | Genes encoding enzymes and their regulators involved in the remodeling of the extracellular matrix |
| **E2F7_E2F1** | E2F transcription factor network |
| **SOX3_SOX2** | Genes encoding structural ECM glycoproteins |
| **HOXB13** | regulation of sodium ion transmembrane transporter activity |
| **MAFF_MAFG** | Genes encoding secreted soluble factors |
| **FOXN1** | Canonical NF-kappaB pathway |
| **FOXD3_FOXI1** | CD40/CD40L signaling |
| **AR_NR3C2** | AP-1 transcription factor network |
| **NFKB1** | CD40/CD40L signaling |
